# Complex Computation from Developmental Priors

**DOI:** 10.1101/2021.03.29.437584

**Authors:** Dániel L. Barabási, Taliesin Beynon, Ádám Katona

## Abstract

Artificial Intelligence (AI) research has provided key insights into the mechanics of learning complex tasks. However, AI models have long overlooked innateness: how strong pressures for survival lead to the encoding of complex behaviors in the nascent wiring of a brain. Although innate neural solutions have inspired AI approaches from layered architectures to ConvNets, the underlying neuroevolutionary search for novel heuristics has not been successfully systematized. In this manuscript, we examine how neurodevelopmental principles can inform the discovery of computational heuristics. We begin by considering the weight matrix of a neural network to be emergent from well-studied rules of neuronal compatibility. Rather than updating the network’s weights directly, we improve task fitness by updating the neurons’ wiring rules, thereby mirroring evolutionary selection on brain development. We find that the resulting framework can not only achieve high performance on standard machine learning tasks, but does so with a fraction of the full network’s parameters. Further, when we condition neuronal identity on biologically-plausible spatial constraints, we discover representations that resemble visual filters and are capable of learning transfer. Finally, we show that developmentally-inspired techniques have higher and more stable performance on metalearning tasks than the standard models they encode. In summary, by introducing realistic developmental considerations into machine learning frameworks, we not only capture the emergence of innate behaviors, but also define a discovery process for structures that promote complex computations.

## INTRODUCTION

The diversity and specificity of animal behaviors, as well as their neural correlates, has received attention from diverse areas of study [1]. Recently, machine learning has provided key insights into the mechanics of solving complex behaviors [2, 3]. However, AI frameworks do not capture the emergence of innate behaviors, as conventional models require extensive update rules and training examples to achieve desired fitness on a task [4, 5]. Nevertheless, a number of complex tasks seem to be hard-coded into the development of the nervous system, such as mice responding to looming stimuli [6], hatched turtles heading out to sea [7], or humans recognizing face-like objects in the womb [8, 9]. In cases where evolutionary pressures for survival outweigh learning, wiring embeddings evolve in order to encode crucial behaviors into the nascent connectome [4]. The brain’s innate solutions have long inspired AI techniques [10], from convolutional neural networks to reinforcement learning, yet evolutionary innovation has not been successfully recapitulated for the systematic discovery of powerful architectures.

In order to reproduce the selection process behind innate behaviors, we must first confront the mystery of the “genomic bottleneck”: development’s uncanny ability to unpack a genome in order to produce specific, task-relevant neural circuits [4, 11]. Although past work has introduced biological considerations for the selection of task knowledge [12, 13], recent studies have mathematically formalized the neuronal partner-selection processes that enable the hard-coding of connectivity [14, 15]. The resulting Genetic Connectome Model considers how interactions between expressed proteins seed synapse formation, and was validated by predicting the wiring rules that code for the *C. elegans* gap junction connectome [15]. Intriguingly, such genetic wiring rules promote structured connectivity, including feed-forward and scale-free networks [16].

In this manuscript we explore realistic developmental mechanisms’ potential for guiding AI exploration. We begin by considering neural network weights to be emergent from the developmental processes formalized in the Genetic Connectome Model [14, 15], thus seeking to learn biologically-validated neuronal wiring rules. By mapping the genetic material passed on in evolution to individual fitness on a task, we provide a mechanistic model for the evolution of innate behaviors. Finally, we examine how incorporating realistic models of cell identity could allow for visual filters to be hard-coded in development. When we test this hypothesis on categorization benchmarks, we find that developmental considerations allow for representations robust enough for both meta- and transfer learning tasks, a hallmark of complex computation.

## GENETIC NEUROEVOLUTION FORMALISM

The processing capabilities of neural systems arise from a mixture of neuronal partner selection, learning, and random noise. Of these, neuronal partner selection, consisting of axonal targeting and synaptic compatibility, provide the basis for hardcoded circuits and innate behaviors. The cellular identity of neurons, as represented by their genetic profile, plays a crucial role in their preferred projections and synaptic partners. This mapping from genes to connectivity is captured by the Genetic Connectome Model [14, 15], which defines the wiring of the brain (**W**) as a function of neuronal identity (**X**) and interactions between genetic factors (**O**). Specifically, the connectivity of *N* neurons is described by the adjacency matrix **W** of size *N*×*N*, with *W_ij_* = 1 if a connection is present between neurons, and 0 otherwise. Individual neurons are identified by their expression of *G* genes, defining a cell-identity matrix **X** of size *N*×*G*, where *X_ia_* = 1 if neuron *i* expresses gene *a* (Figure 1a, blue to orange links). Interactions between expressed genes determine the wiring of the connectome, represented as the *G*×*G* matrix **O** (Figure 1a, orange to orange links). Thus, the developmentally produced connectome can be formulated as

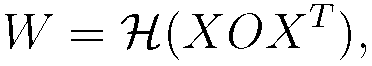

where 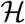 represents the heaviside function that produces a binary connectivity matrix. Previous work used binary values for **W**, **X** and **O** to provide interpretable results for connectivity, genetic interactions and expression patterns [14, 15]. In reality, genes (**X**) have continuous expression, interactions (**O**) may be probabilistic, and connections (**W**) vary in size and number. This prompts a relaxation of **W**, **X**, and **O** to IR, allowing for the three matrices to be continuous and differentiable.

**FIG. 1:**
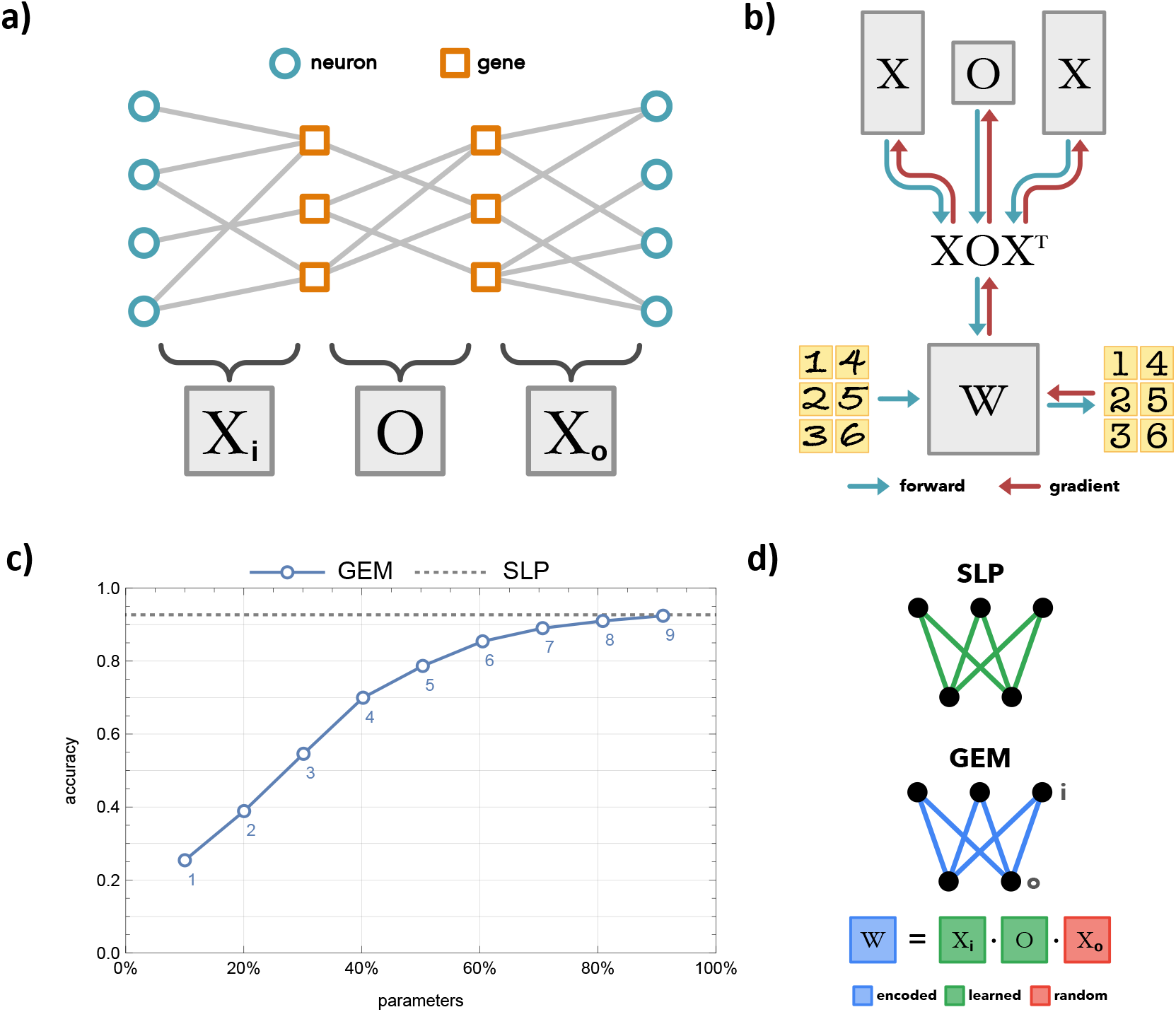
The Genetic neuroEvolution Model. **(a)** Visualization of the Genetic Connectome Model [14, 15]. Matrices *X_i_* and *X_o_* represent the gene expression of input and output neurons, respectively. The **O** matrix corresponds the genetic interactions that underlie neuronal partner selection. **(b)** Traditionally, AI techniques define an architecture (**W**), which can receive data as inputs (handwritten digits, left of **W**), and produce predictions (digital numbers, right of **W**). The weights (**W**) of the architecture can be updated (leftward red arrow) based on the distance of the predictions from known values, thereby producing a more accurate system with training. In the Genetic neuroEvolution Model (GEM), the architecture (**W**) is produced from a small set of wiring rules defined by **X**s and **O** (downward blue arrows). At each training step, the data is passed through to make predictions (rightward blue arrows). However, rather than altering the architecture weights, gradients are computed to update **X**s and **O** (upward red arrows). At the next training step the revised wiring rules generate a revised **W** (downward blue arrows). **(c)** Performance of GEM on the MNIST task. The accuracy of a single-layer linear classifier (784 by 10 nodes) is shown, either with learned weights (dashed line) or weights encoded by GEM’s wiring rules (blue line, with number of genes labeled below each marker). Parameters are expressed as a percentage of a learned SLP’s weights. **(d)** Visualization of a learned SLP and an SLP encoded by GEM in Figure 1c. We do not learn *X_o_*, as we find it adds parameters without increasing task performance.

Based on this extended framework, we propose a Genetic neuroEvolution Model (GEM) that utilizes the generative process of *W* = *XOX^T^* to move flexibly between a wiring diagram and its encoding in neural identities (Figure 1b). We start by taking an architecture known to be effective on a task, then define the weights of the network using 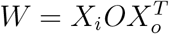 (Figure 1b, downward blue arrows). Here, **W** corresponds to the weights of a layer, *X_i_* stands for the genetic expression of the input neurons, and *X_o_* represents the output neurons’ gene expression. We begin with untrained **X**s and **O** chosen from a fitting random distribution. With each training batch, we backpropagate the loss *through* the weights (Figure 1b, red leftward arrow), updating the **X**s and **O** using gradients calculated by PyTorch’s autograd (Figure 1b, red upward arrows). In theory, this results in a new developmental ruleset in the **X**s and **O**, which produces a more fit **W** in the next task initialization. Thus we adapt a standard AI approach to model the evolution of cell identities and genetic interactions, generating ‘individuals’ that have prewired behavioral programs at ‘birth.’

In order to validate the hypotheses of GEM we turn to the MNIST hand-drawn digit classification task (Figure 1c), which has well-studied accuracies for a number of architectures [17]. We aim to use GEM to learn “developmentally initialized” network weights that have high task performance, without further direct weight updates (“learning”) after initialization. To minimize the number of free variables (i.e. size and number of hidden layers), we start by encoding a single-layer linear classifier (Figure 1d), consisting of 28 ∗ 28 + 10 = 794 nodes (neurons) with 7,840 learnable weights (synapses). A single layer neural network initialized with kaiming uniform performs at an accuracy of 10% without any training (“naive”), however, with direct training of weights can achieve 93% accuracy (“learned”, dashed line in Figure 1c). We aim to encode the weights of this network using GEM, with the number of genes *G* as the only free variable in the system. This results in a parameter scaling of *P* = *N_i_* ∗ *G* + *G* ∗ *G* + *G* ∗ *N_o_* = *G*(*N_i_* + *N_o_* + *G*), where *N_i_* and *N_o_* are the number of input and output neurons, respectively. We find that GEM can achieve nearly 25% accuracy “at birth” when neurons “express” only 1 gene (corresponding to under 10% of the parameters of the full single-layer classifier). The accuracy increases to above 80% for *G* = 5, corresponding to 50% of the parameters of a linear classifier. The GEM model achieves full 93% accuracy with 9 genes, or 90% of the baseline parameters. Thus, we can not only match the task performance of the classically trained network with fewer parameters, but we also achieve high accuracy with half of the original parameters.

## EMERGENCE OF CELL IDENTITY

Current understandings of neurodevelopment indicate that reproducible connectivity arises from interactions between cell identity and neuronal compatibility rules [18]. Thus far, we have used a biologically-validated model of neuronal compatiblity to derive an AI model for the emergence of innate behaviors. Next, we examine how more realistic considerations for cell identity can impact neuroevolutionary search. Although constraining the system could destabilize learning, we aim to show that complex computations can be primed by developmental processes.

To first approximation, a neuron’s cell type (1) defines its connectivity and function, and (2) emerges from well-known factors, such as cell lineage and spatial gradients [19–21]. To make a machine learning analogy, the developmental patterning of cell types corresponds to a reproducible processing hierarchy, which we will assume to be a layered structure (Figure 2a). We can think of these as ‘deterministic’: if the organism develops normally, these cell identities emerge consistently and define the broad strokes of neurons’ projections and connections [22]. However, neurons have also been shown to exhibit a random component to their wiring. For instance, stochastic alternative splicing of cell adhesion molecules allows for a significant amount of genetic variability to emerge [23, 24]. Such processes further partition a single coarse-grained cell type, leading to divergent sub-identities, and thus divergent input and output profiles within a neural population.

**FIG. 2:**
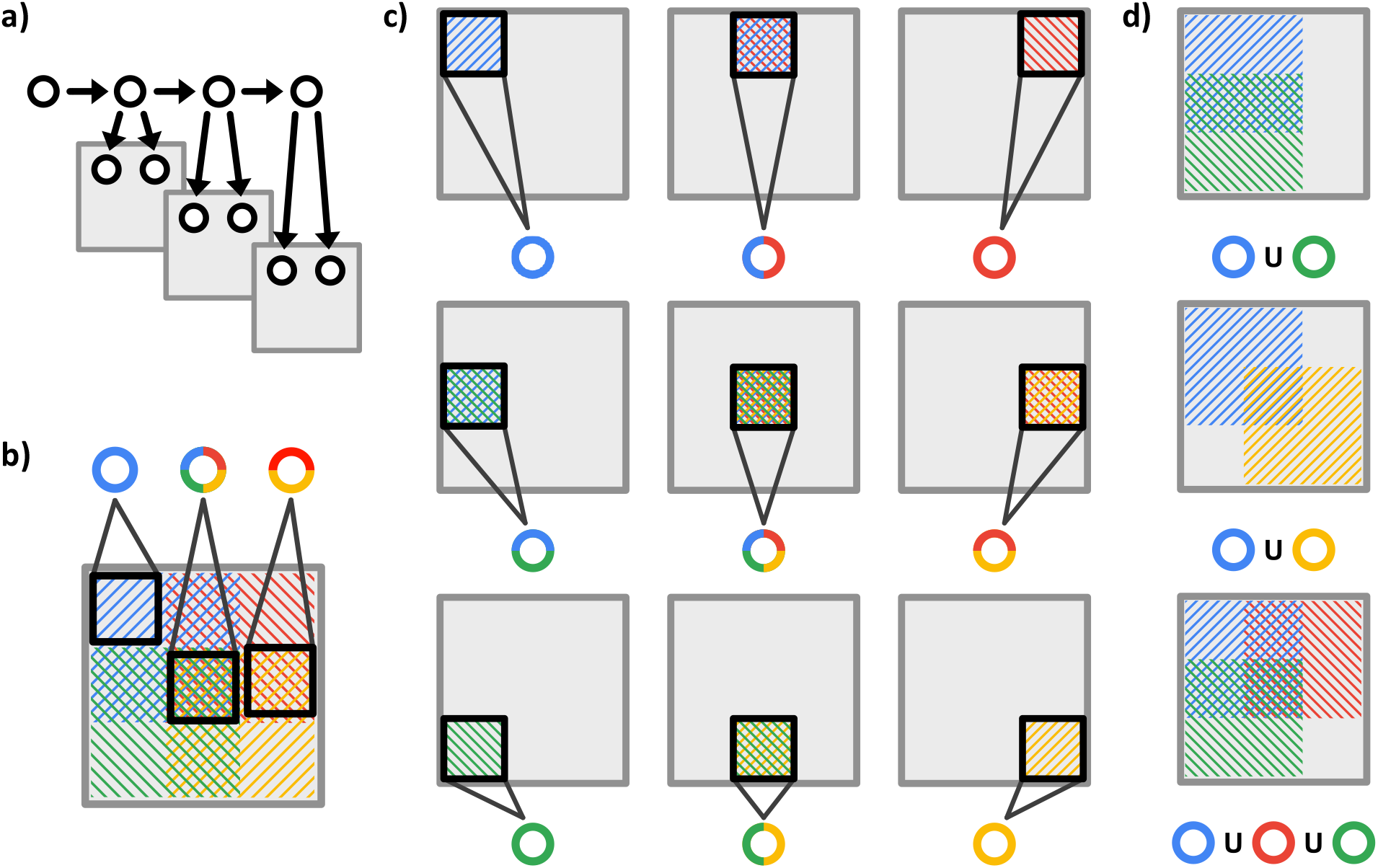
Developmental Priming of Computation. **(a)** Coarse-grained cell types, such as those arising from lineage, can define a neuron’s position in a processing hierarchy. **(b)** Spatial gene expression can produce diversity within the coarse-grained cell types. Here we have four genes (blue, red, green and yellow), each of which has a spatial bias to its expression (shading). When sampling from the top left, we find neurons that express the blue gene (left circle). However, if we sample at locations where the gene expression distributions overlap, we can find neurons with multiple genes expressed, such as red and yellow (right circle), or even all four genes (middle circle). **(c)** Four genes with spatial biases are sufficient for a spatial tiling to be developmentally primed. Neurons can have specific wiring rules, such as connecting to neurons that only expresses blue genes (top left). Other neurons may be more specific, incorporating ‘and’ operators, for instance only receiving connections from neurons expressing both blue and red genes (top, center). **(d)** Incorporating ‘or’ operators allows for the coding of spatial filters. Neurons that allow connections from blue or green genes can select for the left side of the region (top). A diagonal filter can be achieved by selecting for blue or yellow genes (middle). Finally, an upper triangle can be selected by being sensitive to blue, red, or green genes, but not yellow (bottom).

Yet, the emergence of cellular variability is not completely random: the alternative splicing of implicated genes, and thus the resulting cell identity, has crucial biases in space and time [25, 26]. We can represent the expression of genes, or their alternative splices, as a mixture of gaussians, 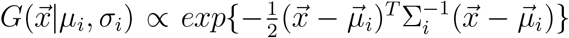, where *G* describes the expression level of a gene over a relevant parameterization of space and time 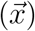. Thus, the spectrum of ‘fine-grained’ cell identities at a given location in the brain is dependent on the distribution of locally expressed genes (Figure 2b). What emerges is a definition of cell types that can be linked not only to a well-studied statistical learning paradigm, but is rooted in specific, highly profiled, genetic mechanisms.

At this point, we have arrived at developmentally motivated models of both cell identity and connectivity rules. As we meld these two components, we find that even simple heuristics applied to a small number of genes with spatial biases can produce complex circuits. Returning to the four genes in Figure 2, we can see that if a neuron only accepts connections from cells expressing blue genes, it will wire to cells in the top left corner of the space (Figure 2c, top left). A different neuron may select for the top center region by accepting connections from neurons expressing *both* blue and red genes (Figure 2c, top middle). As we extend these varying levels of selectivity, we find that we can partition the space in Figure 2c to form a convolutional tiling. Finally, if we incorporate ‘or’ operators, we can produce more specific filters on a given region (Figure 2d). Thus, spatially biased genes and simple wiring rules can begin to reproduce the inductive biases of the convolutional neural networks that underpin modern image recognition solutions.

## SPATIAL PRIORS ON NEUROEVOLUTION

In order to test the hypotheses of Figure 2, we ask whether competitive task performance can be evolved from a GEM approach where cell identity (e.g. **X** matrix) is fully determined from spatial gene expression gradients. We define a spatial GEM by placing N neurons on a grid (28×28 for an MNIST input layer), and determine a neuron’s gene expression profile (**X** matrix) by its distance from *G* 2-D gaussians with given *σ* and *μ* (Figure 3a). As training progresses we fix the locations of neurons and update the gene distributions’ *σ* and *μ*, thereby learning a biologically constrained **X** (Figure 3b). When we use the spatial GEM (S-GEM, Figure 3c right) to encode the SLP from Figure 1c, we achieve over 80% accuracy with less than 2% of the total parameters (Figure 3c, left). S-GEM converges towards 93% accuracy with 10% of total parameters, a point at which the non-spatial GEM only performs at 25% accuracy. To better contextualize the performance of S-GEM, we turn to a random basis encoding (RB, Figure 3c left), which is a measure for the number of free parameters needed to solve a task to a given accuracy [27]. We find that the S-GEM not only matches the parameter-accuracy tradeoff of the random basis model, but outperforms it at low parameter counts.

**FIG. 3:**
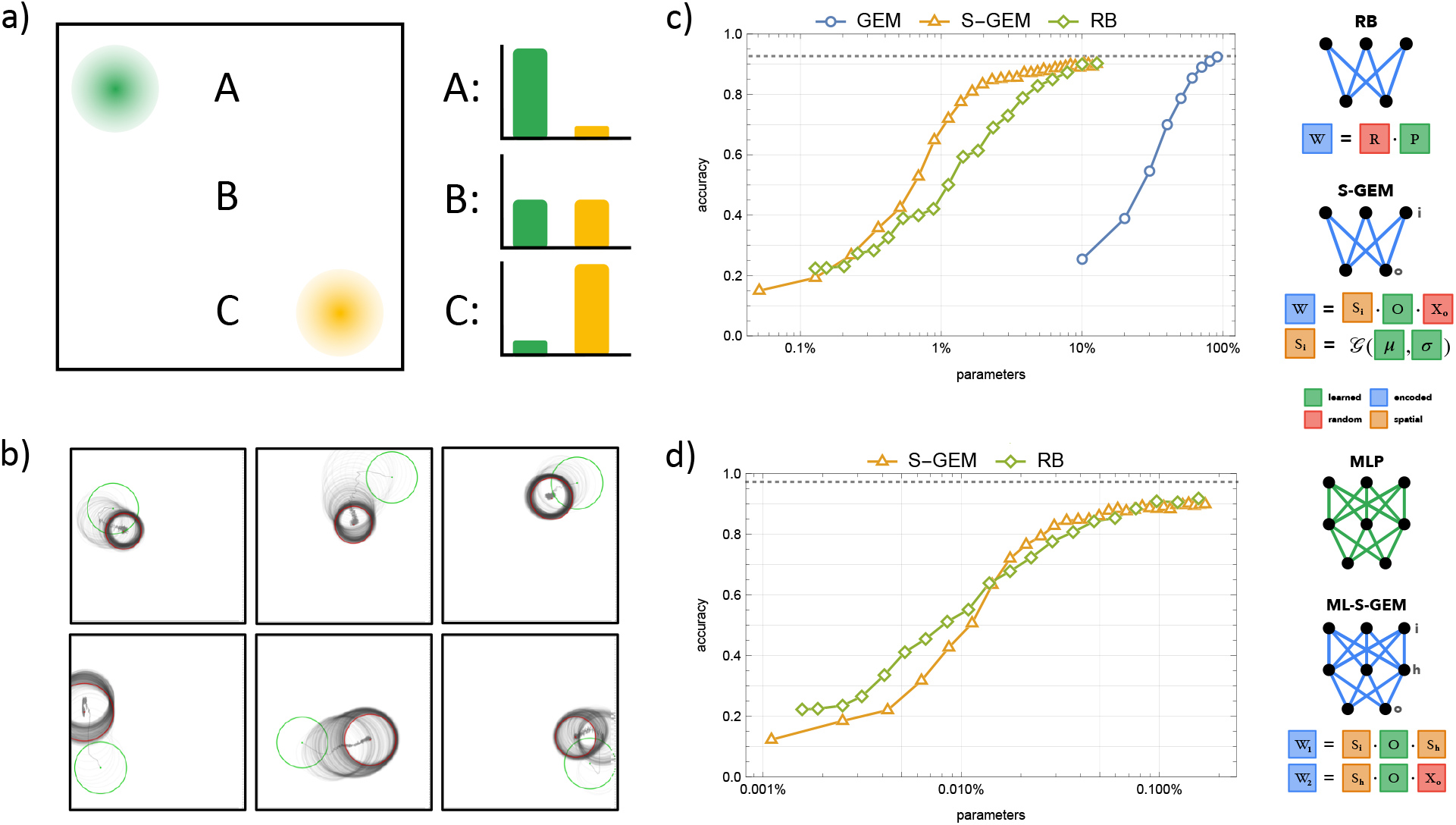
Spatial GEM. **(a)** In the spatial GEM, cell identities are determined by neurons’ distance from 2-D gene expression gaussians. Given two spatial gene distributions, green and yellow, we have: neuron A with high expression of green and low expression of yellow, neuron B with medium expression of both genes, and neuron C with low expression of green but high expression of yellow. **(b)** As learning progresses, the location (*μ*) and dispersion (*σ*) of the gene expression distributions change from start (green) to finish (red). **(c)** Left: Parameter-accuracy tradeoff for encoding a single layer linear classifier that solves MNIST, where parameters are expressed as a % of an uncompressed SLP. The spatial GEM (S-GEM) model significantly outperforms the GEM framework from Figure 1c, as well as a random basis (RB) encoding. Right: Visualization of the used RB and S-GEM weight encodings. **(d)** Left: Parameter-accuracy tradeoff for encoding an MLP that solves MNIST, where parameters are expressed as a % of an uncompressed MLP. The multi-layer S-GEM perform comparatively to a random basis encoding. Right: Visualization of the MLP and ML-S-GEM encoding used.

In order to confirm the robustness of the S-GEM encoding, we extend our validation to multilayer perceptrons (Figure 3d, right). Doing so requires refining our biological assumptions. Given that gene interactions are determined by biophysical rules, and therefore are not specific to a set of neurons, we choose to learn a single **O** matrix that is shared by all encoded layers. Further, we assume that neurons have a single cell identity that determines their input and output connectivity profiles, thus we learn a single **X** matrix for each hidden unit, using it to define both its pre- and post-synaptic weights. Although neurons can express different genes in axons and dendrites, we found that learning separate input and output **X** matrices for each hidden unit only increases the number of parameters, without increasing accuracy. The total parameters of this spatial GEM model scales as *P* = *G* ∗ *G* + *L* ∗ 3*G* = *G*(*G* + 3*L*), as we have a single **O** matrix with *G* ∗ *G* parameters, and *each* encoded layer (L) has a learned *μ_x_, μ_y_* and *σ* for *each* gene. It is interesting to note that the parameter count no longer scales with the number of neurons in the architecture, as cell identities are emergent from the local gene expression gradients. In fact, although we currently restrict neurons to a grid, future work could explore an S-GEM encoding where the locations of neurons are learned, or sampled from a learned distribution, thereby naturally accommodating the addition or subtractions of neurons from a layer.

Given this definition of a multi-layer spatial GEM (ML-S-GEM, Figure 3d right) we aim to encode an MLP of size 784×784×10, where the input and hidden neurons are placed on 28×28 grids. We find that the parameter-accuracy tradeoff continues to match a random basis encoding, achieving above 90% accuracy with less than 0.1% of the encoded MLP’s parameters. As a point of comparison, a less biological “genomic bottleneck” model achieved 79% accuracy with 1038-fold parameter compression on this architecture [11], while the ML-S-GEM achieves above 80% accuracy with roughly 0.025% of the original MLP parameters, amounting to a 4000-fold compression. Even for this larger network, ML-S-GEM trains in seconds to minutes, corresponding to only a slight increase in computational burden compared to a standard MLP. In summary, we find that introducing biologically motivated constraints on cell identity not only retains the performance of our model of innate behavior, but provides additional compression of the hardcoded circuit.

Having confirmed the compression capabilities of S-GEM, we turn to the final prediction of Figure 2: introducing biological biases to gene expression should produce recognizable structures in the networks’ weights. When we observe the weights of a standard MLP trained on MNIST (Figure 4a), we find an uninspiring solution: the hidden nodes seem to learn outlines of numerals, such that the output nodes only have to ‘read out’ the active solutions. In contrast, an MLP encoded by S-GEM exhibits spatial structure in the hidden layer (Figure 4b). We found that this spatial structure was unique to S-GEM, as it does not appear in GEM or Random Basis encodings (Figure S1). Yet, the spatial structure also emerges when S-GEM is used to encode a denoising autoencoder (SI section B, Figures S2 and S3) [28], highlighting the robustness of the phenomenon.

**FIG. 4:**
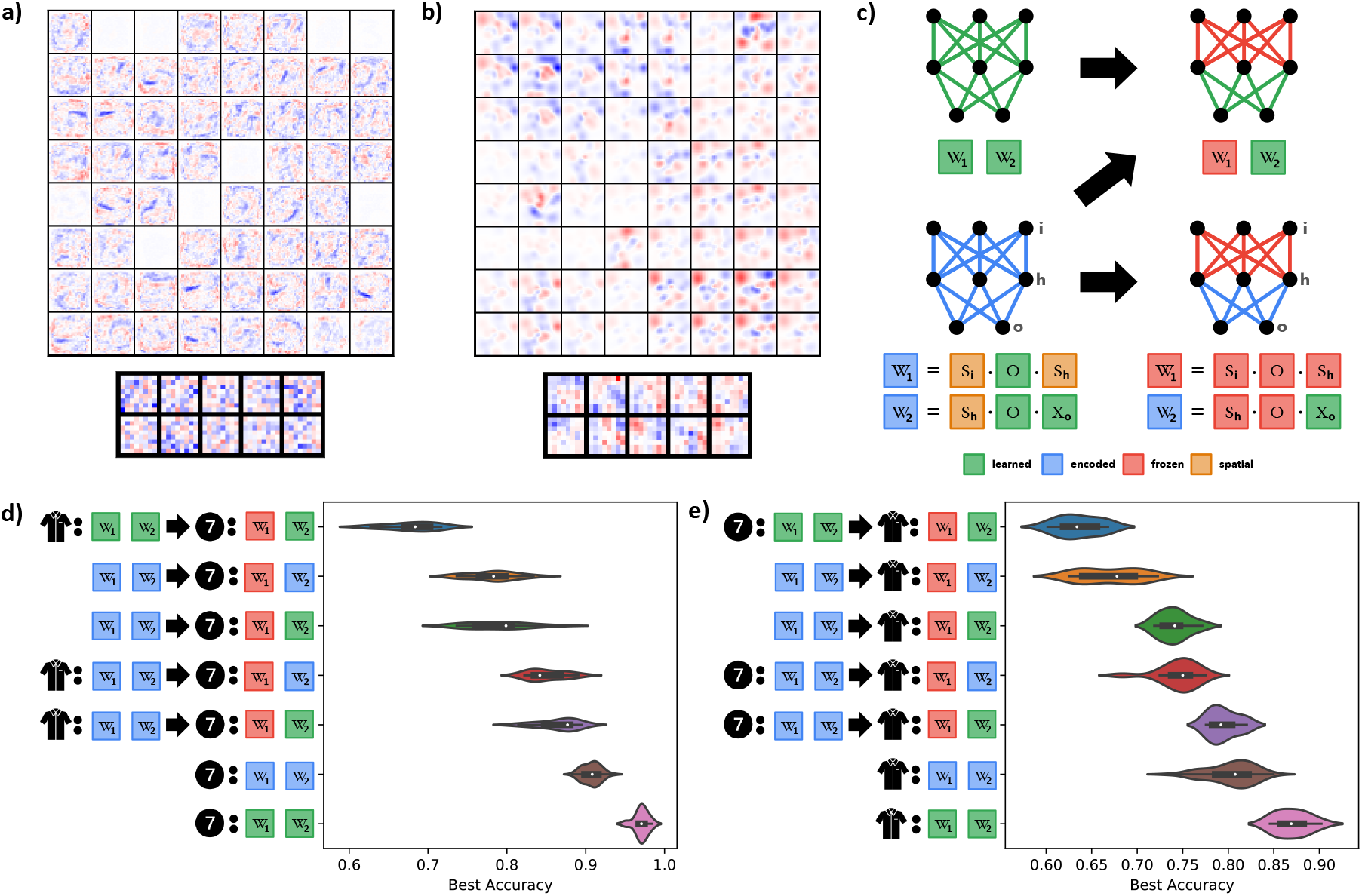
Transfer of Learned Representations. **(a)** Weight matrices for a standard MLP (784×64×10) trained on MNIST. Each square shows the weight matrix of a single node, with inputs and nodes arranged spatially. For instance, the top left node of the hidden layer has weights of the MNIST shape 28×28, where the top left pixel in the box corresponds to the hidden node’s weight from the top left pixel of the input image. **(b)** Weight matrices for an S-GEM (784×62×10) trained on MNIST, laid out as in (a). **(c)** Visualization of transfer learning. A standard MLP (Top, Left) can be trained on one task, e.g. FMNIST, then, with the first layer frozen (Top, Right), can be trained on a second task, e.g. MNIST. Alternatively, an ML-S-GEM model (Bottom, Left) can be trained on a task, then can either be retrained to a new task by updating just the gene expression of the output layer (Bottom, Right) to model “Evolutionary Transfer,” or the output weights can be retrained entirely (Top, Right) to model “Learning Transfer.” **(d)** Violin plots for the performance on MNIST, with or without transfer learning. Top row: MLP is trained on FMNIST (shirt), then retrained on MNIST (number) with the first layer frozen. The next two rows consider architectures with weights initialized by ML-S-GEM, however the first layer was frozen immediately, and the resulting models were trained on MNIST by either learning the gene expression of the output layer (second row) or updating the hidden to output weights directly (third row). The next two rows train ML-S-GEM on FMNIST, and then are trained on MNIST by either learning the gene expression of the output layer (fourth row) or updating the hidden to output weights directly (fifth row). Last two rows contain an ML-S-GEM (sixth row) and standard MLP (last row) trained directly on MNIST. **(e)** Performance on FMNIST, with or without transfer learning. Order of plots rows are akin to (d), however we are now transferring from MNIST to FMNIST.

Given the similarity of the spatial weights of S-GEM to visual filters, we wondered if they contain generalized representations, akin to CNN layers. We tested for generalizability by asking our models to perform transfer learning, which quantifies whether representations learned by a neural network for one task can be efficient on a second task. For instance, we can train an MLP (Figure 4c, top left) on FMNIST, a categorization task for fashion items. Next, we can take the FMNIST-trained MLP and freeze *W*_1_, the weights between the input and hidden nodes (Figure 4c, top right), and train only *W*_2_, the weights between the hidden and output nodes, to solve MNIST. By freezing the first set of weights (*W*_1_), we freeze the way the network represents the FMNIST image in its hidden layer, and therefore ask it to perform on MNIST by learning a readout (*W*_2_) from this constrained “representation space.” If the learned representation is a general computational solution (i.e. visual filters), it should be competitive on the new task. However, a standard MLP (784×784×10) performs poorly on task transfer, achieving less than 70% accuracy on MNIST after training on FMNIST (Figure 4d, top row), compared to the over 90% accuracy of an S-GEM or MLP trained solely on MNIST (Figure 4d, bottom two rows). The results are similar when the order of the tasks is reversed: an MLP trained first on MNIST, then trained with a frozen *W*_1_ on FMNIST performs at under 65% accuracy(Figure 4e, top row), compared to the above 80% accuracy by an S-GEM or MLP trained solely on FMNIST (Figure 4d, bottom two rows).

Given this baseline, we can test the performance of a neural network encoded by S-GEM (Figure 4c, bottom left) on transfer learning. Transfer learning through S-GEM can be done in two ways. One the one hand, we can freeze the spatial gene expression parameters and the **O** matrix, and learn only the gene expression of the output nodes (thereby updating only *W*_2_, Figure 4c, bottom right), which we consider a model of evolutionary adaptations to novel visual environments. Alternatively, can use the weights encoded by S-GEM for *W*_1_, but update *W*_2_ directly through backpropagation, with no encoding (Figure 4c, top right), which we consider on-line transfer learning within an organism’s lifetime. We find that both these approaches significantly outperform the transfer learning of the standard MLP, achieving 80-90% accuracy when going from FMNIST to MNIST (Figure 4d, rows 4 and 5) and 75-80% accuracy when going from MNIST to FMNIST (Figure 4e, rows 4 and 5). As a sanity check, we wanted to ensure that this significant improvement does not come from having introduced relevant spatial information through the genetic gradients. To test this, we initialized networks by S-GEM but did not train them on an initial task before freezing *W*_1_, and found that the resulting random representation (Figure 4d,e rows 2-3) outperformed standard MLP transfer accuracy, but did not achieve equivalent performance to the transferred S-GEM approaches. In summary, we find that the transfer learning capabilities of S-GEM are augmented by pertinent information in the spatial cell identities, but also leverage generalized representations developed through “evolution.”

## DEVELOPMENTAL PRIMING OF LEARNING

Thus far we have focused on modeling evolutionary hardcoded behaviors, where an individual has high fitness at birth. In addition to the prevalence of hardcoded traits, animals also exhibit developmentally primed behaviors, where the evolved connectome promotes the acquisition of a complex task. For instance, humans are not born with the ability to speak, nor are infants’ brains predisposed to learn only their parents’ native tongue. Rather, during a critical period children can rapidly acquire languages they are exposed to. This suggests that development has poised the wiring of language areas to rapidly analyze and distill sound patterns, with local wiring recontouring to encode language in the process. In this section, we show that GEM can evolve condensed wiring rules whose rolled-out nets can flexibly, and rapidly, acquire relevant skills.

We begin by considering the meta-learning topic of few-shot classification [29, 30] as a model of developmental priming of learning. Here, rather than reinforcing high fitness on a task at birth, we aim to select for embeddings that perform well within a few training steps. Developing fit embeddings in this manner can be considered as a nested optimization process: an outer “evolution” loop generates a primed **W** from **X** and **O** while an inner “learning” loop adapts the wiring to the task at hand (Figure 5a).

**FIG. 5:**
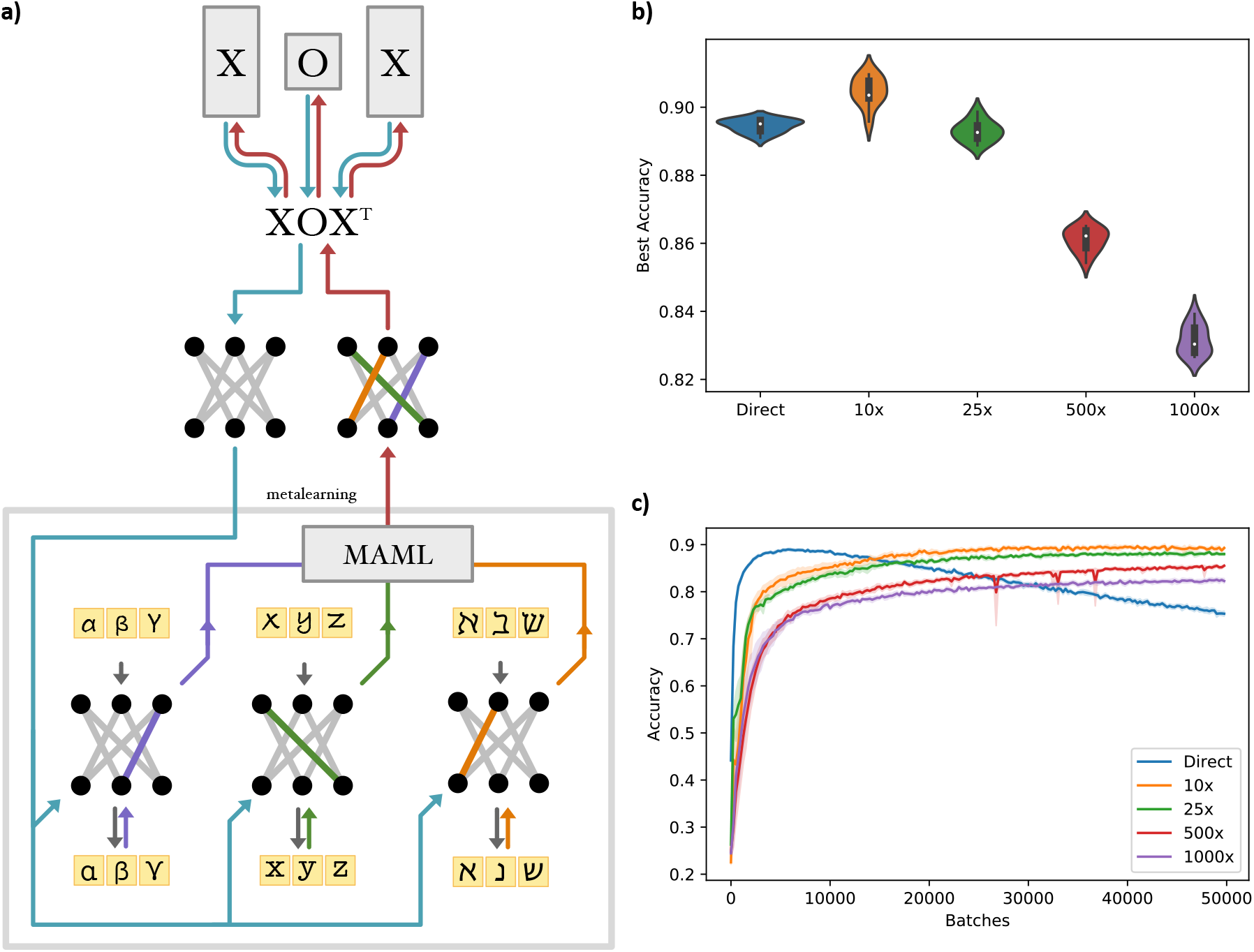
Meta-Learning with GEM. **(a)** Visualization of the Omniglot meta-learning pipeline. In the outer loop (top) we GEM generates an initial network, which is used for all tasks in the inner loop (“metalearning,” bottom). For each classification task (Greek, Latin and Hebrew alphabets) a small number of training steps leads to diverging updates of the model (blue, green and orange weights, respectively). The MAML algorithm allows for a meta-gradient (red arrows) to be passed back to **X** and **O** in the outer loop, thereby generating a network better suited to all tasks in the next iteration. **(b)** Violin plots of the best test accuracy for various models (n = 10 runs per model). In order of performance, a 10x GEM achieves 90.5 ± 0.4% accuracy, a standard MLP (Direct) achieves 89.5 ± 0.2%, 25x GEM achieves 89.3 ± 0.3%, 500x S-GEM achieves 86.1 ± 0.3%, and 1000x S-GEM achieves

Specifically, we turn to the 5-way, 1-shot Omniglot benchmark, which consists of categorizing a library of over 1,000 unique characters from multiple alphabets [31]. We use the MAML algorithm [29] to train linear, feedforward models with two hidden layers of size 784 (see SI C: Meta-Learning on Omniglot). We find that the GEM compression can not only match the performance of a standard MLP (89.5 ± 0.2% best accuracy over 10 runs, Figure 5b “Direct”) with a 25x compression (89.3 ± 0.3%, Figure 5b “25x”), but can outperform it under a 10x parameter compression (90.5 ± 0.4%). We can achieve even higher levels of compression with the S-GEM, however it comes at a slight cost to performance, with a 500x compression achieving 86.1 ± 0.3%, and a 1000x compression achieving 83.2 ± 0.4% accuracy. In addition to this competitive accuracy, the GEM and S-GEM encodings do not overfit to the training data, a major drawback of the standard MLP paradigm (Figure 5c). In summary, we find that developmentally primed encodings outperform traditional ML methods in both accuracy and stability for meta-learning tasks. 83.2 ± 0.4% accuracy. **(c)** Average accuracy of the models over 50,000 training batches (n = 10 runs). Although the standard MLP achieves peak performance in fewer batches, it overfits the training data, decaying in accuracy with further batches. In contrast, both GEM (10x, 25x) and S-GEM (500x, 1000x) compressions achieve peak performance later, but do not suffer overfitting.

## DISCUSSION

In this paper we explored a biologically-validated embedding for the evolution and encoding of innate behaviors. We began from a model of developmental wiring rules, and proposed the Genetic neuroEvolution Model (GEM) in order to “evolve” a neural network that has high task performance “at birth.” Next, we examined how a biologically realistic model of cell identity can promote structured connectivity. When applied to MNIST tasks, we found that incorporating spatially defined genes into the GEM led to a more compact encoding, all while allowing development to prime topographic maps capable of transfer learning. Finally, we showed how utilizing GEM for Omniglot leads to better, and more stable performance when compared to standard MLPs of equivalent size. By understanding, then building on neurodevelopmental embeddings, we expect GEM to provide continued insight for both neuroscience and machine learning.

In defining the spatial GEM we aimed to utilize genetic phenomena profiled in multiple systems and organisms. For instance, stochastic splicing of Pcdh has been implicated in mouse OSN [32], purkinje [32], and cortical microcircuit wiring [33], as well as the somatosensory mapping of the barrel cortex [34]. In *Drosophila* Dscam alternative splicing is crucial for mechanosensory neurons [24]. Perturbation experiments highlight the level of specificity achievable by this genetic mechanism. One method of reducing the available splicing diversity of Dscams in *Drosophila* affected only mechanosensory neurons, which achieved the ‘broad strokes’ of connectivity (initial axon guidance), but lost the specificity of the “last stretch”: where to branch, how many synapses to make, and which neurons to connect to [24]. In mice the Pcdh stochastic expression repertoire was found to be crucial for short-term memory and sensory integration [35].

Given our focus on simple, but well documented, developmental phenomena, further work could benefit from examining genetic and activity-dependent finetunings of neural circuits. For instance, although the topographic mapping from retina to SC is initially established through the graded expression of receptors and ligands [36, 37], the process is aided by interaxon competition [38] and is refined by spontaneous activity before eye opening [36, 37]. Spontaneous activity is also crucial for the alignment of the topographic maps formed from retina to SC and from SC to V1 [39]. Yet, spontaneous activity may play only a guiding role in circuit formation; experiments in ferrets have shown orientation selectivity prior to eye opening, a result which persist even under dark-rearing [40] or silencing of spontaneous activity [41]. Further, ex-vivo preparations, which lack spontaneous activity, still develop proper subcellular projection patterns [42]. These studies suggest that the genetic mechanisms incorporated into the spatial GEM model already play a significant role in the required specificity of hardwired computational circuits.

## Acknowledgements

We wish to thank Gabriel Kreiman, Adam Marblestone, Dániel Czégel, Albert-László Barabási, and the IBRO-Simons Computational Neuroscience Imbizo for fruitful discussions. D.L.B. was supported by NIH NIGMS T32 GM008313 and a Mind Brain and Behavior Graduate Award.

## Author Contributions

Both authors contributed to the design of the project and figures. D.L.B. wrote the manuscript. T.B. developed the code.

## Data and Code Availability

A github repository for the Genetic neuroEvolution Model (GEM) is available at https://github.com/taliesinb/DiffConnect/tree/WIP. The analyses of the study utilized a number of publicly available software packages and datasets which could not be reproduced in the repository, but can be found through in-text citations.

## Conflict of Interest

The authors declare no conflict of interest.

## SI

### A. Learning on MNIST

We implemented MNIST training through the PyTorch deep learning environment. We utilized ADAM with cross entropy loss to train networks for a maximum of 30,000 batches of size 64. We tested the accuracy every 1,000 batches, and included early stopping if the accuracy did not improve by more than 0.005. The accuracy was tested on a hold-out set of 10,000 digits that were not used for training. The accuracy numbers reported in the figures and text are the highest hold-out accuracy that was measured during the training run. Where multiple training runs were involved in producing a single accuracy figure, the mean was taken.

The single layer perceptron (SLP) of Figure 1c was a single linear layer with 784 input and 10 output nodes. We trained all MLPs, encoded or not, with a single 784 hidden layer with ReLU, thereby easily comparing to concurrent models of the genomic bottleneck [11]. The only exceptions were the weights in Figure 4a,b and Figure S1, for which we used a hidden layer of size 8*x*8 = 64 to allow for easier visualizations.

When training the GEM model, we begin with randomly initialized *X_i_*, *X_o_*, and *O* matrices. For each training batch, we generate the weights of a layer using *W* = *X_i_OX_o_*. We evaluate the cross-entropy loss for the batch using these weights, with which we evaluate the gradients for the *Xs* and *O.* We discard the current weight matrix, update *Xs* and *O*, then generate a new weight matrix for the next batch. This is a key difference from the approach of [11], which updates the weight matrix through multiple batches, then learns generating rules that approximate the updated weights. In addition, while [11] alter their parameter count by changing the size of their “g-network” neural network architecture, our parameter count is determined by the number of genes expressed by each neuron. In this way, our parameter count for GEM scales as *P* = *N_i_* ∗ *G* + *G* ∗ *G* + *G* ∗ *N_o_* = *G*(*N_i_* + *N_o_* + *G*), where *N_i_* and *N_o_* are the number of input and output neurons, respectively, and G is the number of genes expressed by a neuron. This means that when G = 3, each neuron is described by a vector of length 3, and an SLP can be encoded by total 2,391 parameters, or roughly 30% of the parameters of an unencoded SLP. At this time we do not include sparsity requirements for *X*s or *O*, although this could provide relevant for further investigations.

Learning with S-GEM proceeds similarly to GEM, with weight generation and developmental rule update in each batch. However, the identities of neurons are no longer a G-length vector learned directly by backprop. Rather, we space the input and hidden neurons evenly on separate 2D grids of size 28×28. Then, we initialize G gene expression gaussians with parameters *σ*, *μ_x_* and *μ_y_*. The gene expression of a neuron j is calculated as 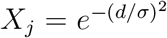, where d is the distance of neuron j from *μ_x_* and *μ_y_*. We introduce two regularizations into this process. First, we add a tanh on the *μ*s, which ensures that we never have gene expressions that are centered outside of the grid of neurons. Second, we apply exp to the variances to ensure that the standard deviations are positive numbers. In each batch we generate the spatial *X*s, which use to form the *W* matrix, and based on the loss update the *σ*s, *μ*s and **O**, thereby learning a biologically constrained **X** (Figure 3b). The total parameters of this spatial GEM model scales as *P* = *G* ∗ *G* + *L* ∗ 3*G* = *G*(*G* + 3*L*), as we have a single **O** matrix with *G* ∗ *G* parameters, and *each* encoded layer (L) has a learned *μ_x_, μ_y_* and *σ* for *each* gene. In all S-GEM instances for MNIST and F-MNIST we do not use a spatial encoding for the output layer, as there is no meaningful way to place 10 nodes on a spatial grid, thus we learn an *X_o_* of size 10xG directly as in GEM. In the case of Figure 3, we simply freeze the randomly generated *X_o_* matrix, as we found learning it only adds additional parameters without increasing accuracy.

For additional implementation details, we refer the reader to the github repository linked in the Data and Code Availability section.

### B. MNIST Autoencoder Task

To test the possibility of hard-coding filters by modeling gene expressions as gaussians, we utilize an MNIST denoising autoencoder task [28]. To begin, we set 10% of pixels in each MNIST digits to 0, and ask a network to reconstruct the original digit. We use a standard architecture [28] of 1 hidden layer with 100 neurons, and input and output layers of size 28 ∗ 28 = 784 neurons (the size of an MNIST image). A sigmoid layer is included after the hidden layer and the output layer. For the spatial GEM approach, we place all the neurons on a grid (10×10 for hidden, 28×28 for input and output), and then pick *G* gaussians, with fixed *σ* but random *μ*, on the grid. This leads to the *NxG*-sized **X** matrix being fixed as the value of the gaussian *g_i_* at that neuron’s location. We then train the **O** matrix, as well as any bias vectors. We compare this to a linear network and a standard GEM approach, where the **X** matrix can be learned. Training occurs on 5000 iterations of batches of size 64. We use a BCE loss, which compares the output image to the initial, unperturbed, MNIST digit.

We find that all three approaches can obtain a BCE of 0.001. However, the linear network requires 157,684 parameters, while the GEM can achieve competitive fitness with 10 genes, corresponding to 17,674 parameters. Finally, the spatial GEM can solve the task with 50 random gaussians, corresponding to only 5,884 trainable parameters.

### C. Meta-Learning on Omniglot

We implemented Omniglot training through the PyTorch deep learning environment. The Omniglot dataset consists of 1623 character classes total from 50 alphabets, of which 1200 classes were used for training, and the rest were reserved for testing [29]. We performed 5-way, 1-shot Omniglot with 2 inner update steps. We utilized ADAM with cross entropy loss and learning rate of 10^−3^ to train networks for a maximum of 50,000 batches of size 32. We learned a unique learning rate for each named parameter, allowing for more specific finetuning as training progressed [43]. We tested the accuracy on a hold-out set every 250 batches, averaging over 10 batches of size 64. The accuracy numbers reported in the figures and text were taken as the mean and accuracy of the highest hold-out accuracy measured in 10 individual training runs.

All trained architectures consisted of a 784 node input layer, two hidden layers of size 784, and an output layer of size 5, with ReLUs interspersed. For the GEM and S-GEM, we learned a single **O** matrix for all layers, but a unique **X** for each layer. The 10x GEM compression corresponded to 50 genes per node, and the 25x GEM used 20 genes per node for the **X** matrices. The 500x and 1000x S-GEM encodings had 43 and 28 spatial genes per layer, respectively.

**FIG. S1:**
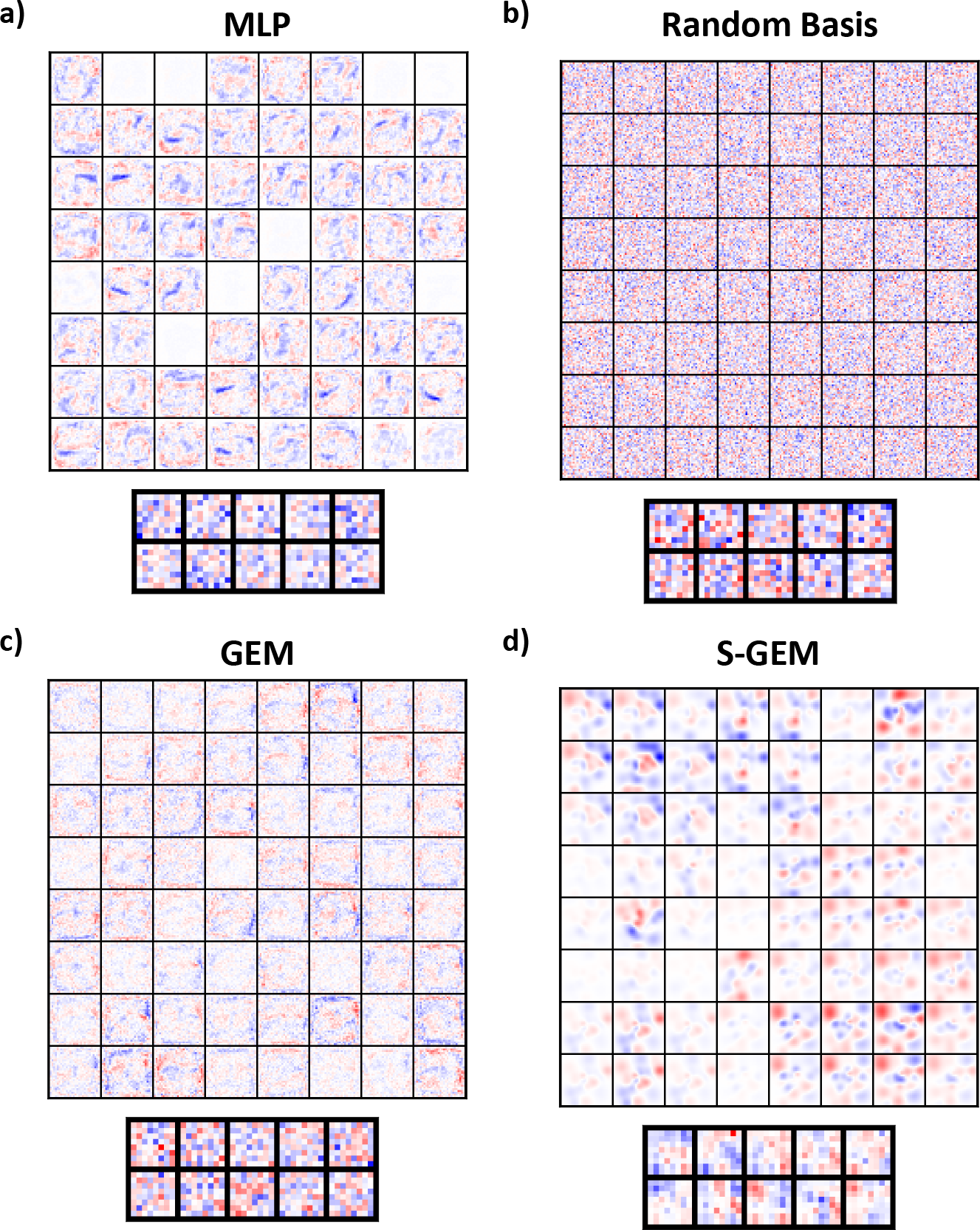
Weight Structure of an MLP Trained on MNIST. **(a)** Weight matrices for a standard MLP (784×64×10) trained on MNIST. Each square shows the weight matrix of a single node, with inputs and nodes arranged spatially. For instance, the top left node of the hidden layer has weights of the MNIST shape 28×28, and the top left pixel in the box corresponds to the top left node in the input layer. **(b-d)** Weight matrices for **(b)** Random Basis, **(c)** GEM, and **(d)** S-GEM encodings of an MLP of size 784×64×10 trained on MNIST, laid out as in (a).

**FIG. S2:**
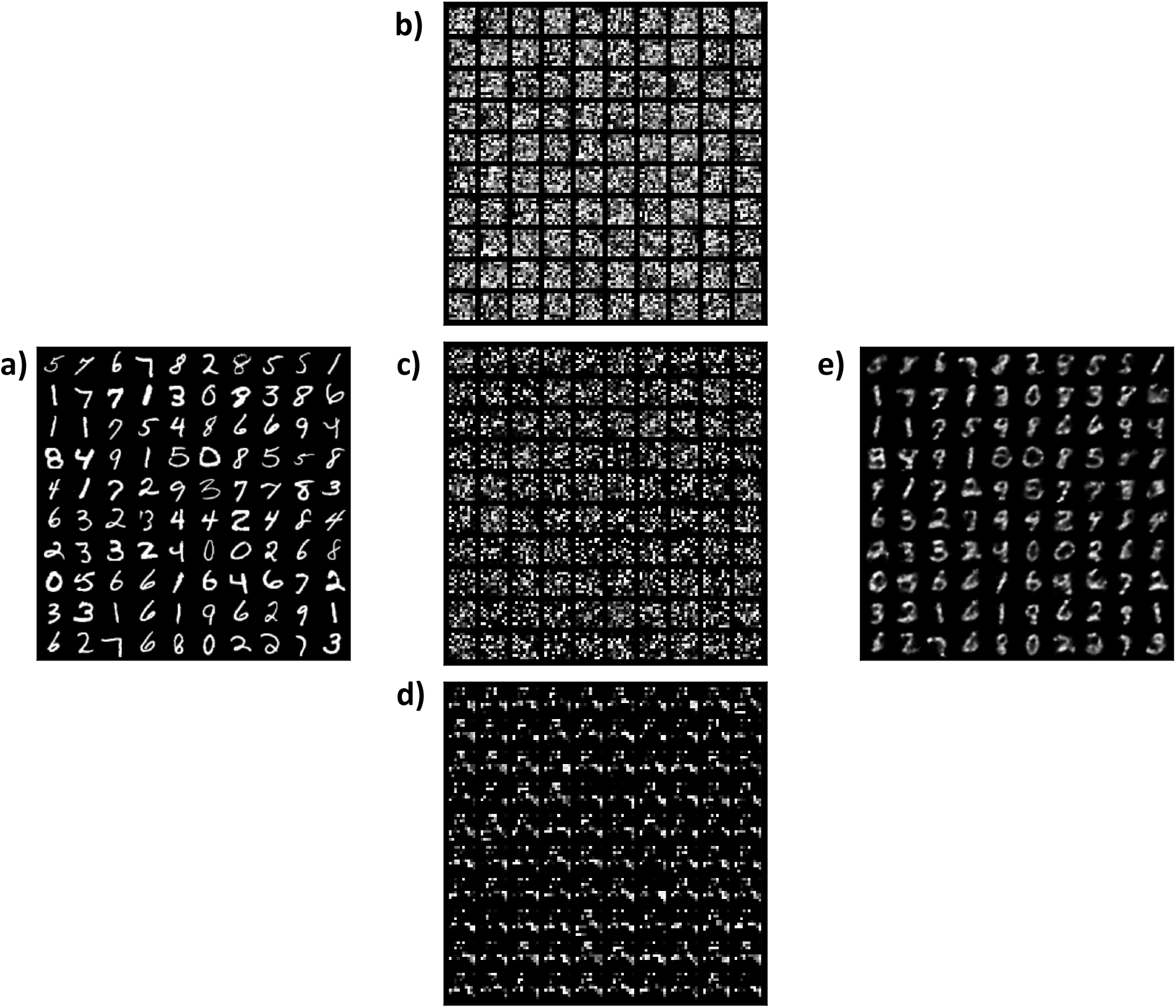
Visual Representation of MNIST Denoising Autoencooder Task. **(a)** Visualization of standard MNIST digits used as input. **(b-d)** Representation of MNIST digits at hidden layer from **(b)** a linear network, **(c)** a standard GEM network, and **(d)** a spatial GEM network. The linear and GEM approaches have noisy, seemingly random, hidden representations, while the spatial GEM approach’s representation recapitulates the spatial structure of the weights (Figure S2). **(e)** Sample output of the task.

**FIG. S3:**
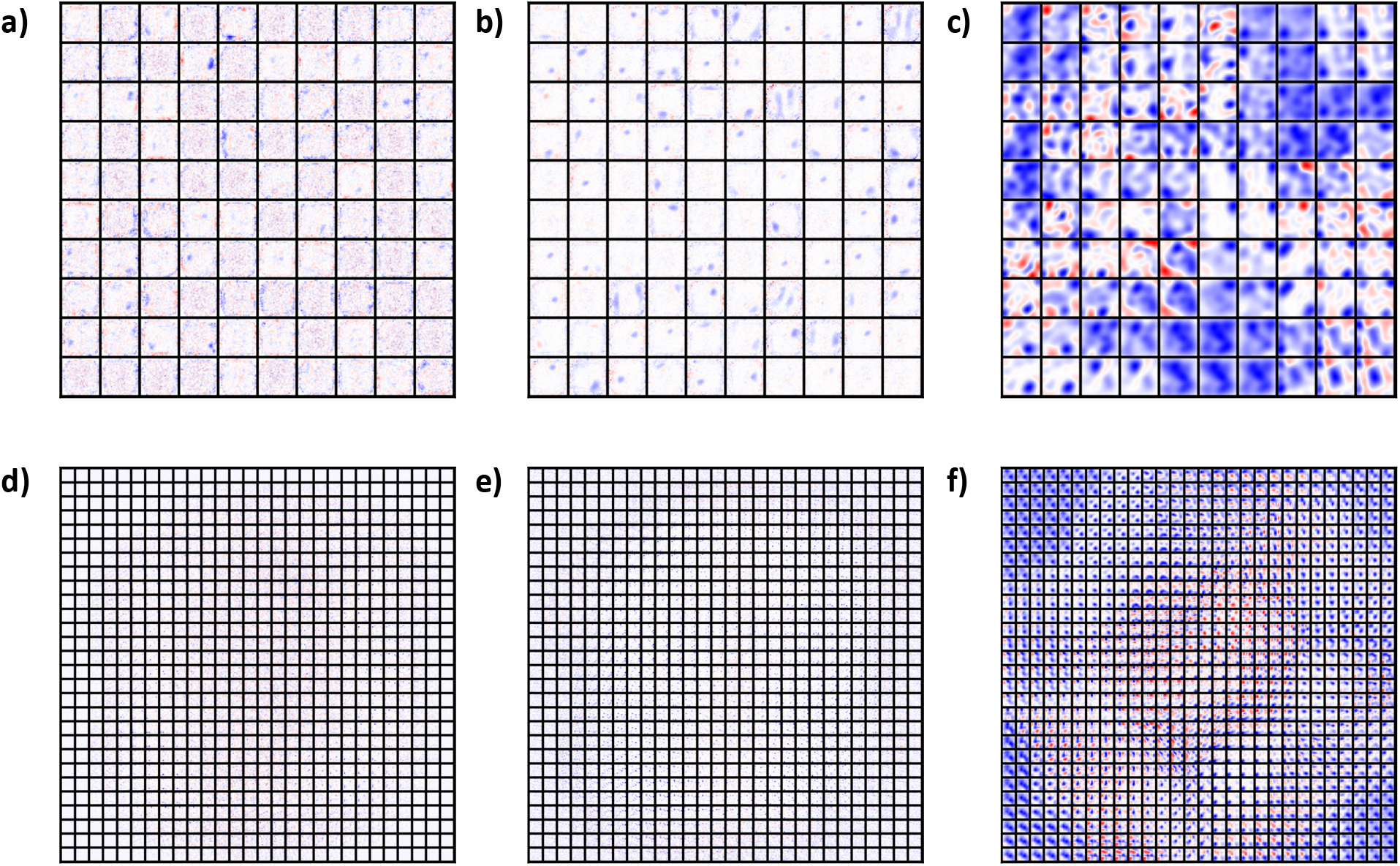
Visualization of MNIST denoising autoencoder weights. **(a-c)** Hidden layer weights for **(a)** a standard linear network, **(b)** a standard GEM network, and **(c)** a spatial GEM network. Red corresponds to positive weights, white is near-zero, and blue is negative. While the spatial GEM network produces weight patterns akin to feature detectors, both the linear and standard GEM approaches learn weights that display less local similarity, both within and between neurons. **(d-f)** Output layer weights of the **(d)** linear, **(e)** GEM, and **(f)** spatial GEM networks. Again, the spatial GEM network displays apparent topographical structure, both within and between neurons, while the other two approaches learned heterogeneous weights.

## Notes

https://github.com/taliesinb/DiffConnect/tree/WIP

### Competing Interest Statement

The authors have declared no competing interest.

### Summary of Updates

Additional figures and authors.

## References

[1] S. Musall, A. E. Urai, D. Sussillo, and A. K. Churchland, Harnessing behavioral diversity to understand neural computations for cognition, Current opinion in neurobiology 58, 229 (2019).

[2] B. A. Richards, T. P. Lillicrap, P. Beaudoin, Y. Bengio, R. Bogacz, A. Christensen, C. Clopath, R. P. Costa, A. de Berker, S. Ganguli, et al., A deep learning framework for neuroscience, Nature neuroscience 22, 1761 (2019).

[3] S. Srinivasan, R. J. Greenspan, C. F. Stevens, and D. Grover, Deep (er) learning, Journal of Neuroscience 38, 7365 (2018).

[4] A. M. Zador, A critique of pure learning and what artificial neural networks can learn from animal brains, Nature communications 10, 1 (2019).

[5] T. P. Lillicrap and K. P. Kording, What does it mean to understand a neural network?, arXiv preprint arXiv:1907.06374 (2019).

[6] M. Yilmaz and M. Meister, Rapid innate defensive responses of mice to looming visual stimuli, Current Biology 10.1016/j.cub.2013.08.015 (2013).

[7] A. F. Carr and L. H. Ogren, The ecology and migrations of sea turtles. 4, the green turtle in the caribbean sea., Bulletin of the AMNH (1960).

[8] V. M. Reid, K. Dunn, R. J. Young, J. Amu, T. Donovan, and N. Reissland, The Human Fetus Preferentially Engages with Face-like Visual Stimuli, Current Biology 10.1016/j.cub.2017.05.044 (2017).

[9] N. Reissland, R. Wood, J. Einbeck, and A. Lane, Effects of maternal mental health on fetal visual preference for face-like compared to non-face like light stimulation, Early Human Development 151, 105227 (2020).

[10] D. Hassabis, D. Kumaran, C. Summerfield, and M. Botvinick, Neuroscience-inspired artificial intelligence, Neuron 95, 245 (2017).

[11] A. Koulakov, S. Shuvaev, and A. Zador, Encoding innate ability through a genomic bottleneck, bioRxiv (2021).

[12] D. Ha, A. Dai, and Q. V. Le, Hypernetworks, arXiv preprint arXiv:1609.09106 (2016).

[13] K. O. Stanley, D. B. D’Ambrosio, and J. Gauci, A hypercube-based encoding for evolving large-scale neural networks, Artificial life 15, 185 (2009).

[14] D. L. Barabási and A.-L. Barabási, A genetic model of the connectome, Neuron 105, 435 (2020).

[15] I. A. Kovács, D. L. Barabási, and A.-L. Barabási, Uncovering the genetic blueprint of the c. elegans nervous system, Proceedings of the National Academy of Sciences 117, 33570 (2020).

[16] D. L. Barabasi and D. Czegel, Constructing graphs from genetic encodings, bioRxiv (2020).

[17] Y. LeCun, L. Bottou, Y. Bengio, P. Haffner, et al., Gradient-based learning applied to document recognition, Proceedings of the IEEE 86, 2278 (1998).

[18] T. C. Südhof, Towards an understanding of synapse formation, Neuron 100, 276 (2018).

[19] S. N. Sansom and F. J. Livesey, Gradients in the brain: the control of the development of form and function in the cerebral cortex, Cold Spring Harbor Perspectives in Biology 1, a002519 (2009).

[20] G. Fishell and A. Kepecs, Interneuron types as attractors and controllers, Annual review of neuroscience 43(2019).

[21] G. Ciceri, N. Dehorter, I. Sols, Z. J. Huang, M. Maravall, and O. Marín, Lineage-specific laminar organization of cortical gabaergic interneurons, Nature neuroscience 16, 1199 (2013).

[22] A. S. Bates, J. Janssens, G. S. Jefferis, and S. Aerts, Neuronal cell types in the fly: single-cell anatomy meets single-cell genomics, Current opinion in neurobiology 56, 125 (2019).

[23] B. Treutlein, O. Gokce, S. R. Quake, and T. C. Südhof, Cartography of neurexin alternative splicing mapped by single-molecule long-read mrna sequencing, Proceedings of the National Academy of Sciences 111, E1291 (2014).

[24] B. E. Chen, M. Kondo, A. Garnier, F. L. Watson, R. Püettmann-Holgado, D. R. Lamar, and D. Schmucker, The molecular diversity of dscam is functionally required for neuronal wiring specificity in drosophila, Cell 125, 607 (2006).

[25] S. L. Zipursky and J. R. Sanes, Chemoaffinity revisited: dscams, protocadherins, and neural circuit assembly, Cell 143, 343 (2010).

[26] G. Neves, J. Zucker, M. Daly, and A. Chess, Stochastic yet biased expression of multiple dscam splice variants by individual cells, Nature genetics 36, 240 (2004).

[27] C. Li, H. Farkhoor, R. Liu, and J. Yosinski, Measuring the intrinsic dimension of objective landscapes, arXiv preprint arXiv:1804.08838 (2018).

[28] C. Fernando, D. Banarse, M. Reynolds, F. Besse, D. Pfau, M. Jaderberg, M. Lanctot, and D. Wierstra, Convolution by evolution: Differentiable pattern producing networks, in Proceedings of the Genetic and Evolutionary Computation Conference 2016 (2016) pp. 109–116.

[29] C. Finn, P. Abbeel, and S. Levine, Model-agnostic meta-learning for fast adaptation of deep networks, in Proceedings of the 34th International Conference on Machine Learning-Volume 70 (JMLR.org, 2017) pp. 1126–1135.

[30] A. Nichol, J. Achiam, and J. Schulman, On first-order meta-learning algorithms, arXiv preprint arXiv:1803.02999 (2018).

[31] B. Lake, R. Salakhutdinov, J. Gross, and J. Tenenbaum, One shot learning of simple visual concepts, in Proceedings of the annual meeting of the cognitive science society, Vol. 33 (2011).

[32] D. Canzio and T. Maniatis, The generation of a protocadherin cell-surface recognition code for neural circuit assembly, Current Opinion in Neurobiology 59, 213 (2019).

[33] X. Lv, S.-Q. Ren, X.-J. Zhang, Z. Shen, T. Ghosh, A. Xianyu, P. Gao, Z. Li, S. Lin, Y. Yu, et al., Tbr2 coordinates neurogenesis expansion and precise microcircuit organization via protocadherin 19 in the mammalian cortex, Nature communications 10, 1 (2019).

[34] T. Hirayama, E. Tarusawa, Y. Yoshimura, N. Galjart, and T. Yagi, Ctcf is required for neural development and stochastic expression of clustered pcdh genes in neurons, Cell reports 2, 345 (2012).

[35] T. Yamagishi, K. Yoshitake, D. Kamatani, K. Watanabe, H. Tsukano, R. Hishida, K. Takahashi, S. Takahashi, A. Horii, T. Yagi, et al., Molecular diversity of clustered protocadherin-*α* required for sensory integration and short-term memory in mice, Scientific reports 8, 1 (2018).

[36] D. Tsigankov and A. A. Koulakov, Sperry versus hebb: Topographic mapping in isl2/epha3 mutant mice, BMC neuroscience 11, 1 (2010).

[37] A. D. Huberman, M. B. Feller, and B. Chapman, Mechanisms underlying development of visual maps and receptive fields, Annu. Rev. Neurosci. 31, 479 (2008).

[38] J. W. Triplett, C. Pfeiffenberger, J. Yamada, B. K. Stafford, N. T. Sweeney, A. M. Litke, A. Sher, A. A. Koulakov, and D. A. Feldheim, Competition is a driving force in topographic mapping, Proceedings of the National Academy of Sciences 108, 19060 (2011).

[39] J. W. Triplett, M. T. Owens, J. Yamada, G. Lemke, J. Cang, M. P. Stryker, and D. A. Feldheim, Retinal input instructs alignment of visual topographic maps, Cell 139, 175 (2009).

[40] L. E. White, D. M. Coppola, and D. Fitzpatrick, The contribution of sensory experience to the maturation of orientation selectivity in ferret visual cortex, Nature 411, 1049 (2001).

[41] B. Chapman and M. P. Stryker, Development of orientation selectivity in ferret visual cortex and effects of deprivation, Journal of Neuroscience 13, 5251 (1993).

[42] G. Di Cristo, C. Wu, B. Chattopadhyaya, F. Ango, G. Knott, E. Welker, K. Svoboda, and Z. J. Huang, Subcellular domain-restricted gabaergic innervation in primary visual cortex in the absence of sensory and thalamic inputs, Nature neuroscience 7, 1184 (2004).

[43] A. Antoniou, H. Edwards, and A. Storkey, How to train your maml, arXiv preprint arXiv:1810.09502 (2018).

